# Validated ligand geometries for macromolecular refinement restraints and molecular mechanics force fields

**DOI:** 10.1101/2025.08.01.668229

**Authors:** Nigel W. Moriarty, David A. Case, Dorothee Liebschner, Paul D. Adams

## Abstract

In macromolecular structure refinement the low observation-to-parameter ratio and the lack of high-resolution data is countered by using *a priori* information in the form of restraints. Having accurate geometries of the chemical entities in the sample is paramount for generating accurate chemical restraints and, therefore, accurate macromolecular structures. In particular, it is desirable to have accurate restraints for known and novel ligand entities. Quantum Mechanics (QM) can minimise the energy of a ligand by adjusting its geometry, and these geometries can be used to generate restraints macromolecular refinement. We describe here a library of 37,000 small molecules extracted from the Chemical Components Dictionary in the Protein Data Bank and minimized by density functional QM. The library includes restraint files for use in crystallography or cryo-EM refinement, along with files suitable for molecular dynamics simulation. Because the geometries are validated, the restraints library provides users with both functional restraints and minimised geometries. This work also provides procedures for generating new and accurate restraints.

## Introduction

Refinement of macromolecular models is driven by both the experimental data and restraints. One universally applied form of restraints are geometry restraints, which provide *a priori* chemical information about the structural components of macromolecules. This chemical information about components is expressed with ideal (or target) values for bonds, valence angles, torsion angles, planes and chirality (Evans, 2007), along with estimated standard deviation (e.s.d.) values that represent the uncertainty of these targets. All refinement programs rely on restraints to produce chemically reasonable models. Restraints for common entities are stored in libraries that are used by most of the refinement programs. Commonly the Engh and Huber library (Engh & Huber, 1991, 2006, 2012) is used for amino acids, and target values from Gilski et al. (2019) and Clowney et al. (1996) for nucleic acids. For example, both phenix.refine (Afonine *et al*., 2012) and REFMAC (Murshudov et al., 2011), use the Engh and Huber restraints or derivatives of them.

Macromolecular structures generally contain protein and/or RNA/DNA. Both of these are polymers described by the restraints libraries as individual sets of restraints for each of the amino acids and nucleic acids. The polymerisation procedure in each program applies restraints to connect each unit in the chain. Another polymer class are carbohydrates, which also have units with restraints and a procedure to apply the appropriate links. Non-polymer entities include ions (mostly single metal ions coordinated to other parts of the macromolecule), metal clusters (also coordinating), and ligands (generally non-covalently bound molecules). Metals often adopt particular geometries with coordinating residues or form clusters with other metals and waters which benefit from restraints in refinement (Moriarty & Adams, 2019; Bhowmick *et al*., 2023). However, it is algorithmically challenging to determine bonds and angles involving metals, and their coordination is further limited by a fundamental shortfall of the overarching restraint formalisation (Moriarty & Adams, 2019). Therefore, metals and metal containing entities are not the scope of this work.

The main topic of this work is ligands but non-standard amino acids are also considered. For the purposes of this study, ligands are molecules that contain at least two atoms and are not classified as polymers or ions^1^. As of April 2025, more than 75% models in the Protein Data Bank (PDB) (Burley et al., 2019; Berman et al., 2003; wwPDB consortium, 2019; Berman et al., 2007) contain at least one ligand. Restraints for ligands can be obtained in two ways: restraints of known ligands can be found in libraries (e.g. HEM and ATP), and those of novel ligands can be obtained from dictionary generators such as ACEDrg (Long *et al*., 2017), GRADE (Smart *et al*., 2011) or eLBOW (Moriarty *et al*., 2009).

Although conceptually similar, libraries of ligand restraints face challenges compared to libraries of amino acids or nucleic acids. First, there are many more ligands than standard (or common non-standard) amino acids. For example, the PDB currently contains over 42,000 non-polymer entities (ligands) corresponding to over 90% of the Chemical Component Dictionary/Library (CCD) (Westbrook *et al*., 2015) entries (the remaining 10% correspond to polymerisable entities or metal clusters; the ligand percentage will only increase as non-polymer additions will outpace polymer additions by a wide margin). Second, the composition of a ligand library needs to be constantly updated. That is, with each deposited model that contains a novel ligand, a new entry needs to be added to the ligand library. Adding restraints for new components is complicated by the fact that the PDB does not currently provide the restraints for the entities, only the bonding information and coordinates, and these do not necessarily represent ideal or representative geometries of the molecule. Third, the development of new therapeutics is exploring new regions of chemical space. This leads to never before seen moieties that can cause issues with any restraints generation technique. Last, many ligands occur only in a single instance of a PDB entry. Indeed, approximately 73% of the CCD entries only occur in a single deposited model, as opposed to standard amino acids (GLY with nearly 230,000 PDB entries), ions (MG, ∼25k), carbohydrates (NAG, ∼11k), nucleic acids (U, ∼8k) and the most common ligand – HEM (∼6k instances). Even for those restraints that occur in less than a few PDB entries, it is desirable that the generation be as robust and accurate as possible.

The Monomer Library (Vagin *et al*., 2004; Murshudov *et al*., 2011) is a restraints library that contains restraints for amino acids, nucleic acids, carbohydrates and common ligands. Versions of it have been used by several structural biology software packages, such as REFMAC in the CCP4 suite, in the Phenix suite (Liebschner *et al*., 2019) and TNT (Tronrud *et al*., 1987) in Global Phasing. Each program uses the ideal and e.s.d. values to construct a complete internal representation of the restraints. For example, the program will lookup the ideal bond length between two atoms of a model based on their residue and atom names. This means that every instance of this bond, such as the Cβ–CƔ bond in an arginine residue, will have the same ideal value. Polymerisation of the amino acids, nucleic acids and carbohydrates is performed algorithmically using other information contained in the library.

Phenix originally adopted a trimmed version of the Monomer Library that retained a minimal set of restraints including those for amino acids, nucleic acids, small molecules and carbohydrates. Other most prevalent ligands were also included. The reason for retaining only a limited number of entries in the library was that the torsion restraints, either the ideal values, e.s.d. values or the periodicity, were often not sufficiently validated. Torsion restraints describe the conformations of a ligand, or puckers. As opposed to existing refinement programs, where torsion restraints are not used by default, Phenix does apply these restraints, so it was necessary to adopt a library where these restraints are as correct as possible. The lack of validated torsion restraints applied particularly to carbohydrates but also nucleic acids and, to a lesser extent, amino acids (for example the H atom in NH2 groups).

Therefore, we introduced a new restraints library in Phenix known as the Geometry Standard (GeoStd) to house updated versions of the restraints. The GeoStd started with improved restraints for the standard amino acids that included the Rotamer Library (Lovell *et al*., 2000) and updates of it (Hintze *et al*., 2016). Ideal values for both amino acids and nucleic acids were adjusted as were the restraints constructs such as non-periodic torsion values. This involves the addition of a feature in the torsion restraints that allowed the listing of ideal torsion values rather than relying solely on the periodicity feature. This is particularly important in rotamers as not all of the periodic wells are actually minima. An example is the chi2 dihedral of tryptophan which exists in the 0°, 90° and -90° configurations but not 180°. This scenario cannot be described with periodicity, so the capability to allow discrete values is an advantage. Other examples of improvements of the GeoStd are the inclusion of arginine restraints that accounted for the asymmetry of the side-chain and the deviation of the C`ẟ` atom of the guanidinium moiety (Moriarty, Liebschner *et al*., 2020); iron-sulphur cluster restraints and linkage to protein improvement (Moriarty & Adams, 2019); protonation specific restraints for Histidine (Moriarty, 2024); and using QM to generate accurate geometries for radicals (Liu *et al*., 2022). Recent improvements of the Engh & Huber values (Engh & Huber, 2012) are also implemented (Moriarty & Adams, 2021). The GeoStd initially contained just over 500 entries. This work expands the library to 38,140 entries.

Another example of updated restraints is the conformation dependent library (CDL, (Moriarty, Tronrud *et al*., 2014; Moriarty, Adams *et al*., 2014; Moriarty *et al*., 2016), which allows backbone bond lengths and angles to vary with backbone conformation. The nuance with this approach is that the ideal and e.s.d. values are applied dynamically, dependent on the local geometry of each residue. Instead of applying a single target value from the standard restraints libraries, the values vary with the ɸ and Ѱ torsion angles of the protein backbone. Phenix has also recently added an implementation of the nucleic acid geometry conformation dependent library RestraintLib (Kowiel *et al*., 2016, 2020; Gilski *et al*., 2019) – another configuration dependent library that includes functionals for more fine-grained values.

All the entities (i.e. amino acids, nucleic acids, ligands, etc) in the Protein Data Bank are contained in the CCD under the title – “The chemical component dictionary: complete descriptions of constituent molecules in experimentally determined 3D macromolecules in the Protein Data Bank”. While each entry is stored in CIF format (Crystallographic Information Format (Brown & McMahon, 2002)) it does not contain restraints. Among other information, the entries contain up to two sets of Cartesian coordinates (the first instance of deposited coordinates as well as an ideal calculated geometry), atom names, a bond list and several SMILES specifications (Weininger, 1988). Other fields include the entity name, ligand code, entity parent and date updated to name a few.

Ideally, a general ligand restraints library would provide restraints for all entities in the CCD. This would make it straightforward to refine any model deposited in the PDB, without the need to generate restraints. However, as the content of the CCD is constantly updated, such a library would require constant updates as well. It is unrealistic that such updates can be maintained manually. On the other hand, it is challenging to automatically compute accurate restraints for new or updated ligands. The vast chemical space can result in moieties that have not been encountered before, including novel element combinations, protonation, charges and radicals. QM Minimisation relies on correct charge, protonation and radical state is a requirement, and if these requirements are not met, the minimisation can converge poorly or fail, and the resulting geometries can be poor or wrong. A simple example of inaccurate geometry arises at an acid moiety in Figure 1. If the acid is not protonated (left side) in the experiment, but protonated in the QM calculation (right side), the resulting bond lengths and bond angles will be inaccurate. For these reasons, ligands restraints had been added *ad hoc* to the GeoStd. Many of the *ad hoc* entries were calculated with QM methods and manually validated in various ways.

**Figure 1.**
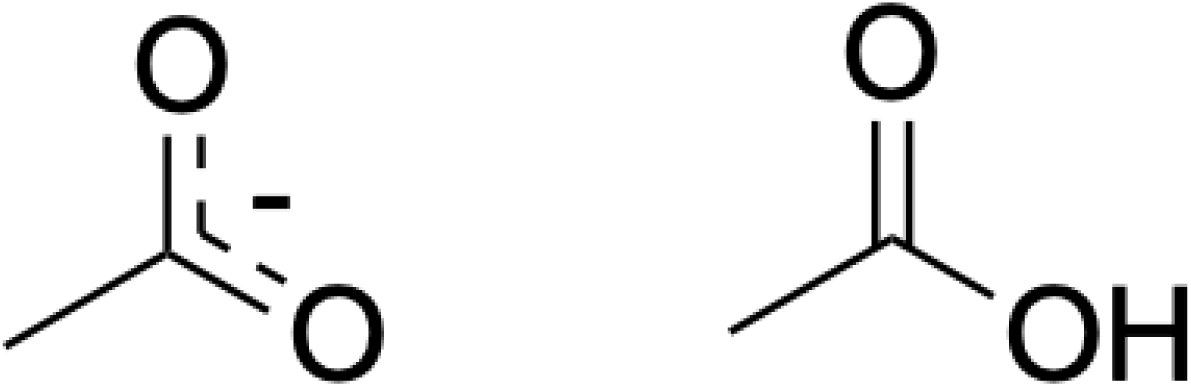
Example of an unprotonated (left) and protonated (right) acid showing acetate and acetic acid.

As the generation of ligand restraints is a common bottleneck for users of crystallographic software, we decided to significantly increase the number of ligand entries in the GeoStd, and to standardise and streamline the addition of novel restraints. This work describes the procedure used to add more ligand entries to the GeoStd restraints library. We describe how entries to be added were selected, how the restraints were generated and how the minimised geometries were validated.

## Methods

To add new entries in the GeoStd, we first determined suitable targets from the CCD (section xxx). We then generated restraints via quantum mechanical methods (section xxx). The minimised geometries were then validated (section xxx). Finally, we also added the functionality of creating force field files for the AMBER molecular mechanics software (section xxx).

### Calculations

All polymer and non-polymer entries in PDB entries are contained in the CCD. As described above, the information stored in the files in this database does not correspond to that present in restraint files, although the file format (CIF) is the same. We used the following information stored in these CIF files: The canonical SMILES of the molecule, the list of atoms, and the list of bonds. Specifically, the list of atoms defines their names which are used to match ligand atoms in a model with the corresponding restraints in a library. Each CCD entry has a second list of legacy atom names if they have been updated since the creation of the initial entry. This may occur during periodic remediation operations performed by the PDB, such as one effort focused on carbohydrates (Access Improved Carbohydrate Data at the PDB) and another effort in 2007 that changed the Hydrogen atom names of amino acids (known as v2 to v3 name conversion).

The version of CCD on 30 July 2024 contained 44,095 entities, including all amino acids, nucleotides, saccharides, ions, metal clusters and non-polymers (ligands).

As a first step, entities in the CCD were filtered. We excluded standard amino acids and nucleic acids, as their restraints are already established and validated. Furthermore, we ignored single atom entries (mostly metal ions, or inert gases, e.g. PDB ID: 7q38) because metal coordination is not in the scope of the GeoStd. We also filtered ligands incorporating metals or metal clusters (e.g., HEM or SF4). To perform QM calculations, metal entries require details of the metal charge and multiplicity (electron distribution in the orbitals). The multiplicity is not provided by the CCD and the charge often changes upon interaction with receptors, so the restraints would not be generally applicable. Furthermore, higher levels of QM are required for accurate computations involving metal bonding.

In addition, we ignored entries that were already in the GeoStd and that had passed previously applied validation. We also did not reprocess manually edited entries, such as the radical MTN for which the plane restraint was adjusted using REEL (Moriarty *et al*., 2017) to account for the added flexibility of the radical moiety (Liu *et al*., 2022) or CLA, for which different planarity restraints for low, medium and high resolution regimes were created and contributed by Dale Tronrud. Any CCD entries denoted as obsolete were also ignored.

To update restraints efficiently, we also checked if a CCD entry had changed significantly since its initial addition to the GeoStd. As an example, when the atom names or the SMILES of an entry in the CCD had been updated, they were accordingly updated in the GeoStd entry. If the molecule content in the CCD was changed compared to what was present in the GeoStd, the corresponding entry was recreated by the procedure described below.

Our work on a new implementation of joint X-ray neutron refinement (Liebschner *et al*., 2019) highlighted the need for supporting different versions of restraints for a ligand, i.e., supporting all possible protonation states of PDB entities. By default, eLBOW generates restraints for typical pH levels in the protein environment. This means, for example, unprotonated acid moieties. However, protonated acid groups are often detected in neutron refinement. The pipeline used in this work to generate the restraints determines if a moiety may be protonated and computes all minimized geometries and corresponding restraints. Accordingly, the restraints lookup in Phenix has been changed to load the appropriate restraints based on the model contents.

After filtering, the majority of remaining components are non-polymer molecules, for which it is generally easier to calculate minimised geometries as they are not covalently bound to other entities. For these components, the environment can be approximated by the use of solvent models and the generation of Amber force fields is simplified.

Some additional steps are needed for polymerisable components. For example, non-standard amino acids in the CCD have neutral moieties on the C and N terminals which is not the situation in the midst of protein (peptide plane) or at the termini of the protein chain (charged). The neutral entity is helpful for QM calculation convergence and a semblance of chemical realism but not a representative geometry of the sample. The restraints are adjusted as discussed in the Restraints Generation section.

### QM Methods

After filtering the CCD entries to obtain ligands to add or update in the GeoStd, the geometry of each molecule in the subset was minimized using quantum mechanical methods.

Many QM methods exist to predict the behavior of molecules, and the methods have varying levels of sophistication. The challenge is that the choice of the method and level of sophistication need to balance the required accuracy of the result (such as the minimized geometry of a molecule) and the computational cost. The entity geometries were minimised using a series of two QM methods: PBEh-3c (Grimme *et al*., 2015) with the CPCM solvent treatment (Barone & Cossi, 1998) and B3LYP D3BJ TightSCF RIJCOSX def2/J def2-TZVP (also known as B3LYP/TZVP). If the PBEh-3c method failed validation, the second method was used. The PBEh-3c method is considered a “high-speed” method but with 17 hours on a modern MacBook Pro, the time to minimise ATP is greater than the average user is willing to wait. Both sets of calculations were performed using the ORCA software package (Neese *et al*., 2020). A semiempirical QM method – PM6-D3H4 (Řezáč *et al*., 2009) – was also assessed for suitability due to its computational efficiency and ease of use via Mopac (Moussa & Stewart, 2024) as distributed with Phenix.

### Restraints Generation

Once the minimized molecular geometries of the ligands were obtained with QM methods, the eLBOW program was to generate restraints. This allowed for adjusting the dihedral restraints type and redundancy. In particular, the restraints generation mechanism in eLBOW ensures that all hydrogen atom restraints are complete. This is relevant for refinement, because phenix.refine uses a riding hydrogen model (Liebschner et al., 2020) to maintain the ideal positions of hydrogen atoms during refinement. The method relies on bond lengths, bond angles and torsion restraints to position the H atom.

Some entities, such as standard amino acids, primarily exist in polymerised form. Polymerisation changes the chemical details of the entities so the restraints need to be modified compared to the single entity versions.. While standard amino acids were not reprocessed in this work, polymerisation can also occur for other entities, such as non-standard amino acids. QM minimisation was performed on the neutral entity as it is present in the CCD, but, to obtain appropriate targets that facilitate automatic polymerisation in Phenix, the superfluous atoms at the termini involved in polymerisation (H2, OXT and HXT) were removed from the restraints. We note that if such a polymerisable moiety occurs at a terminal, correct termination (such as adding back the H2 group) can be handled algorithmically in Phenix.

We note that the PDB recently moved to allow describing ligand IDs with five characters (instead of three) to take into account the growing number of polymer entities (Bank). With Phenix tools being able to process these entries, our approach was also applied to ligands with 5-character ID. Accordingly, the resulting new version of GeoStd includes 505 components with 5-character ID.

### Validation

The program Mogul (Bruno *et al*., 2002; Macrae *et al*., 2008) in the Cambridge Structure Database suite (CSD) (Groom *et al*., 2016) allows the validation of the ligand geometries. It uses accurate experimentally determined small molecule geometries from CSD entries to provide ideal and standard deviation (s.d.) values for bonds and angles which can be used to validate the QM geometries before being added to the GeoStd.

To validate the ligand geometries based on the Mogul s.d. values, we used Z-scores. The Z-score measures by how many standard deviations a specific metric (a bond length, a bond angle, etc) deviates from its target value. For example, an angle Z-score of 1.0 means that a particular angle deviates by one standard deviation. We note that the PDB validates ligands using the Z-scores of the bonds and angles using the Mogul s.d. values. The ligand taskforce (Adams *et al*., 2016) suggested that the absolute root mean squared Z-score (r.m.s.Z) be less than 2 while the PDB report calculates both the r.m.s.Z and displays the number of bond/angle Z-score greater than 2. Experience has shown that the s.d. values from Mogul are generally too small for practical purposes as e.s.d. values in refinement at resolutions typically achieved in macromolecular crystallography. It is more appropriate to use twice the experimental s.d. values from Mogul for e.s.d. values in refinement (Moriarty & Adams, 2019; Moriarty, Liebschner *et al*., 2020; Moriarty, Tronrud *et al*., 2014).

We adopted the following validation scheme using the maximum Z-score. If the maximum (absolute) Z-score was less than 2, 4 or 6, the geometry was designated as “perfect”, “grand” and “ok”, respectively.

As noted above, Mogul generally provides small s.d. values if there are only a few moieties of one type. Further, an extreme scenario occurs when there is only one example of a specific bond or angle in the CSD. This can result in an s.d. value of zero and validation failure. To include such geometries, *ad hoc* minimum s.d. values for bond and angle of 0.005Å and 0.75°, respectively, are applied. These values are quite conservative as they are approximately half the normal s.d. values. If the validation passes using these *ad hoc* s.d. values the geometries are designated “ok (reasonable std)” (no matter the Z-score). Another special case exists for polymerisable entities. The polymerisation of the amino acids means that the ideal values of bonds and angles in the polymer backbone are different from the isolated entity. It follows that the bond and angle ideal values can be updated in the restraints based on the parent type of the amino acid without the need for validation. However, validation is required for the side chain. A geometry validation that is side chain specific has “(side chain)” added to the designation.

### Force Field files

Macromolecular refinement results can be improved using molecular mechanics force fields such as those in Amber (Case *et al*., 2023; Moriarty, Janowski *et al*., 2020). Accurate non-polymer geometries can be used to generate Amber force fields to streamline Amber based refinement. While a tool exists in Phenix to generate force field parameters based on a model the presence of curated force fields aids in speed and accuracy of Amber refinement. In addition, the Amber force fields can also be used in other calculations such as molecular mechanics and molecular dynamics.

For each ligand structure, we used the *antechamber* and *parmchk2* program in AmberTools (Case *et al*., 2023) to prepare force field files using the general Amber force field (GAFF2) (Wang *et al*., 2004) and the ABCG2 atomic charge model (Sun *et al*., 2023; He *et al*., 2025). Since the input geometries had already been minimized at a QM level, minimization was not performed prior to the charge calculation. We also filtered out entities that GAFF2 cannot handle: elements other than H, C, N, O, F, Cl and I; components that are parts of polymers; and the “low-pH” entries with protonated carboxylic acids. This led to force field files for 37,788 components. Each component was then minimized in the resulting force field, to assess how far the force-field optimized geometry deviates from the input, QM-optimized geometry. This was performed using the `gb=1` generalized Born solvent model and a monovalent salt concentration of 0.1 M.

Although the AmberTools programs were used to generate the force field files, the results can be used in many other molecular simulation programs. The *ParmEd* facility in AmberTools facilitates transfer to other common simulation programs such as OpenMM (Eastman *et al*., 2024), GROMACS (Páll *et al*., 2020) and CHARMM (Brooks *et al*., 2009).

## Results

Of the 44,095 entries in the CCD, 42,150 remained after filtering as starting points for QM minimisations. QM minimisation at the PBEh-3c level resulted in 40,342 optimised geometries. Of these, 32,849 passed validation (81%) as seen in the “PBEh-3c” column of Table 1, the majority with the “grand” designation. The “perfect” and “ok” categories also have a large number of entries. We note that a significant number of components were assigned to the “reasonable std” category indicating that many Mogul s.d. values were too small. We also note that the “side chain” classification is a small fraction of the total.

**Table 1.** Number of validation results in each category of the PBEh-3c (both protonated (pH=0) and unprotonated (pH=7) and B3LYP/TZVP calculated molecules.

| Validation class |  | PBEh-3c | B3LYP/TZVP | PBEh-3c (pH=0) |
| --- | --- | --- | --- | --- |
| PERFECT |  | 5566 | 1723 | 2801 |
| PERFECT | side chain | 64 |  | 16 |
| GRAND |  | 15315 | 552 | 1171 |
| GRAND | side chain | 791 | 3 | 97 |
| OK |  | 6363 | 1186 | 1546 |
| OK | reasonable std | 4750 | 1369 | 451 |
| TOTAL |  | 32849 | 4833 | 6082 |

The second QM level was applied to the 16% (6,381) of filtered entries that had not passed the first round of validation. This time, 4,833 (76%) entries passed the validation with the majority of entries having either “perfect” or “ok” geometry. We note that once again the application of *ad hoc* s.d. values added a significant number of entries. The set of calculations is restricted to the failed entries from the previous QM level which may explain the reduced success rate (76% vs 84%). In total 37,682 restraints files were added to the GeoStd using the validated QM geometries.

As noted, restraints were also calculated for protonated states of the ligands, denoted pH=0 in Table 1. This procedure checked whether a protonated state exists for a ligand before expending resources. It was also only performed using the PBEh-3c method. The table shows that 6,082 ligands have protonated restraints that passed validation with the majority having a “perfect” classification.

The size of the molecules has a non-linear impact on the efficiency of the calculations. Even the simplest QM methods scale as the cube of the number of atoms, with more complex QM scaling at higher powers. Many techniques are employed to lower the scaling but they generally use a reasonable interaction cutoff for larger molecules. The size of entities calculated by the PBEh-3c QM method are displayed in Figure 2 with the raw data in Table S1. The information is binned in groups of ten non-hydrogen atoms. The majority of molecules are smaller than 60 atoms which falls within the 20-70 range as delineated by Ghose *et al*. (Ghose *et al*., 1999) for the typical molecule size of a drug candidate.

**Figure 2.**
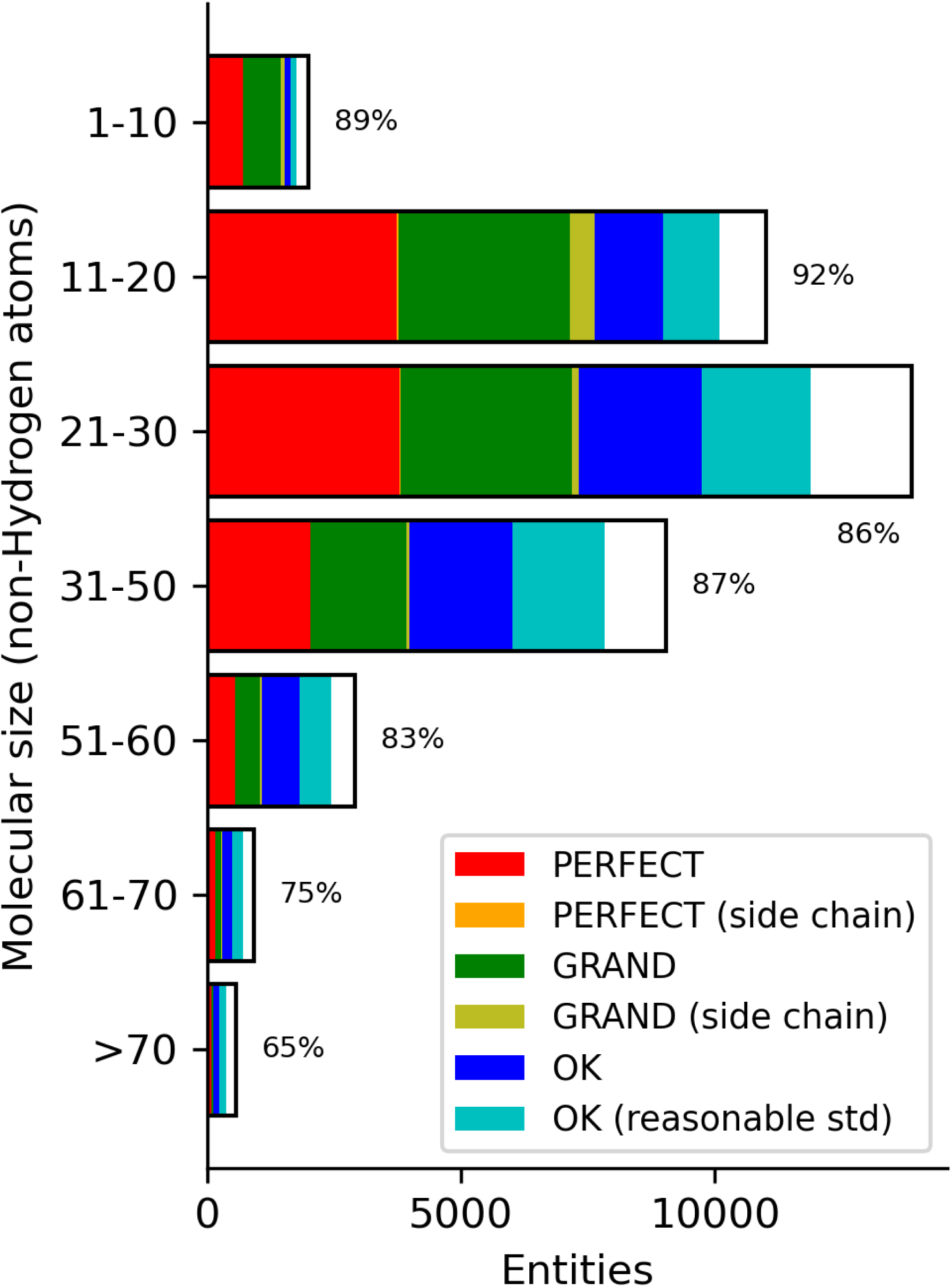
Number of PBEh-3c geometries calculated for each molecular size binned in width of 10. Colours in the bar indicate the number of validation classifications per the legend with the white portion representing the fail classification. The total success rate is displayed next to each bar.

The proportion of the validation categories are shown as different colours in the bar with the white portion indicating those geometries that failed Mogul validation. The majority of geometries in the druglike size regime pass Mogul validation with the smaller molecules being more likely to be accurate. Smaller molecules (1-10) had 29% in the “perfect” classification while large components (>70 atoms) had only 12%. We observe a similar behaviour for the “grand” classification. Unsurprisingly, the “ok” classifications increase as a fraction of the total as the molecular weight increases (6% for 1-10 increasing to 20% for 41-50). The “reasonable std” group is about the same as the unmodified e.s.d. group in all molecular size bins.

Specific moieties are another characteristic of a molecule that influences the outcome of a QM calculation. This can be investigated by the molecular charge and element composition. The distribution of the molecular charge of the PBEh-3c geometries is shown in Figure 3 (raw data in Table S2). Unsurprisingly, the majority of the entities are neutral as is the policy of the CCD. However, because the process used to generate restraints automatically deprotonates acid moieties, there is also a significant number of negatively charged ligands.

**Figure 3.**
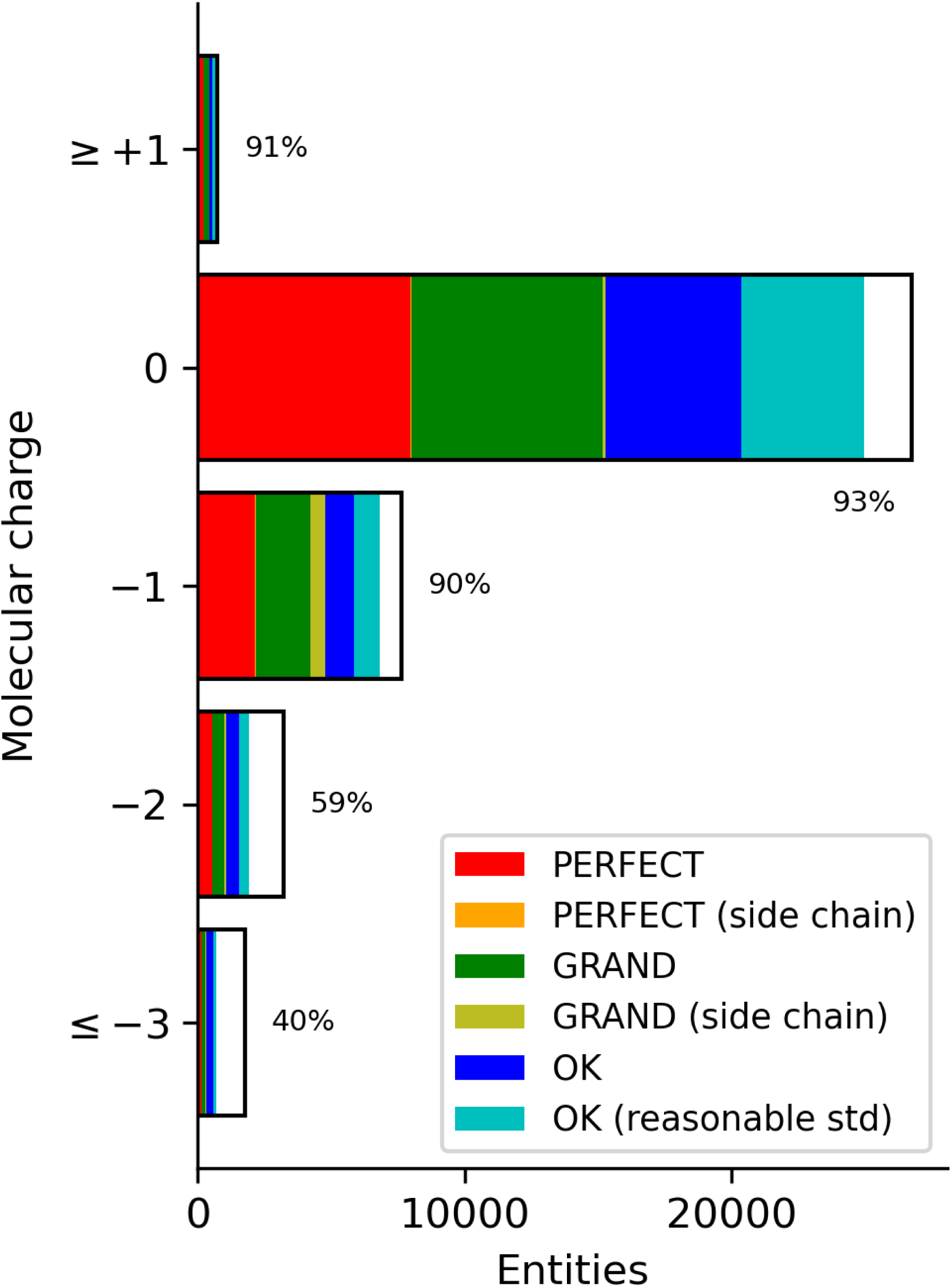
Number of calculated molecules in charge. Colours in the bar indicate the number of validation classifications per the legend with the white portion representing the fail classification.

The validation success rate of the singly charged molecules is nearly 90% while neutral and positively charged molecules are in the 70% range. More negatively charged molecules have lower success rates.

The element content of a molecule can affect the validation outcome for a calculated geometry. It is also a proxy for moieties that can be difficult to minimise to a geometry that will pass validation. Figure 4 shows the validation success rate based on the element content of the molecule. Molecules with halogens had the highest success rate (Br – 92%, Cl – 92%, I – 92% and F – 90%). Note that error bars are included based on population with some bars being too small to be displayed. Molecules containing carbon, nitrogen and oxygen atoms (the bulk of the molecules) have a rate of approximately 87%. The lowest significant success rate was for molecules containing Phosphorus atoms at 41%. These molecules are of particular note because the Phosphorus is often in a phosphate moiety – the charge on the oxygen atoms causing problems for the QM calculations. Alternatively, it may be because many phosphates in the PDB and CSD are coordinated with a Magnesium ion. The presence of the ion greatly affects the bond lengths and angles of the coordinating molecule that are not reflected in the QM calculations of the isolated entity. The presence of selenium in a molecule also appears to present some problems. However, there are only about 120 Se containing ligands in the CCD so the impact is more limited.

**Figure 4.**
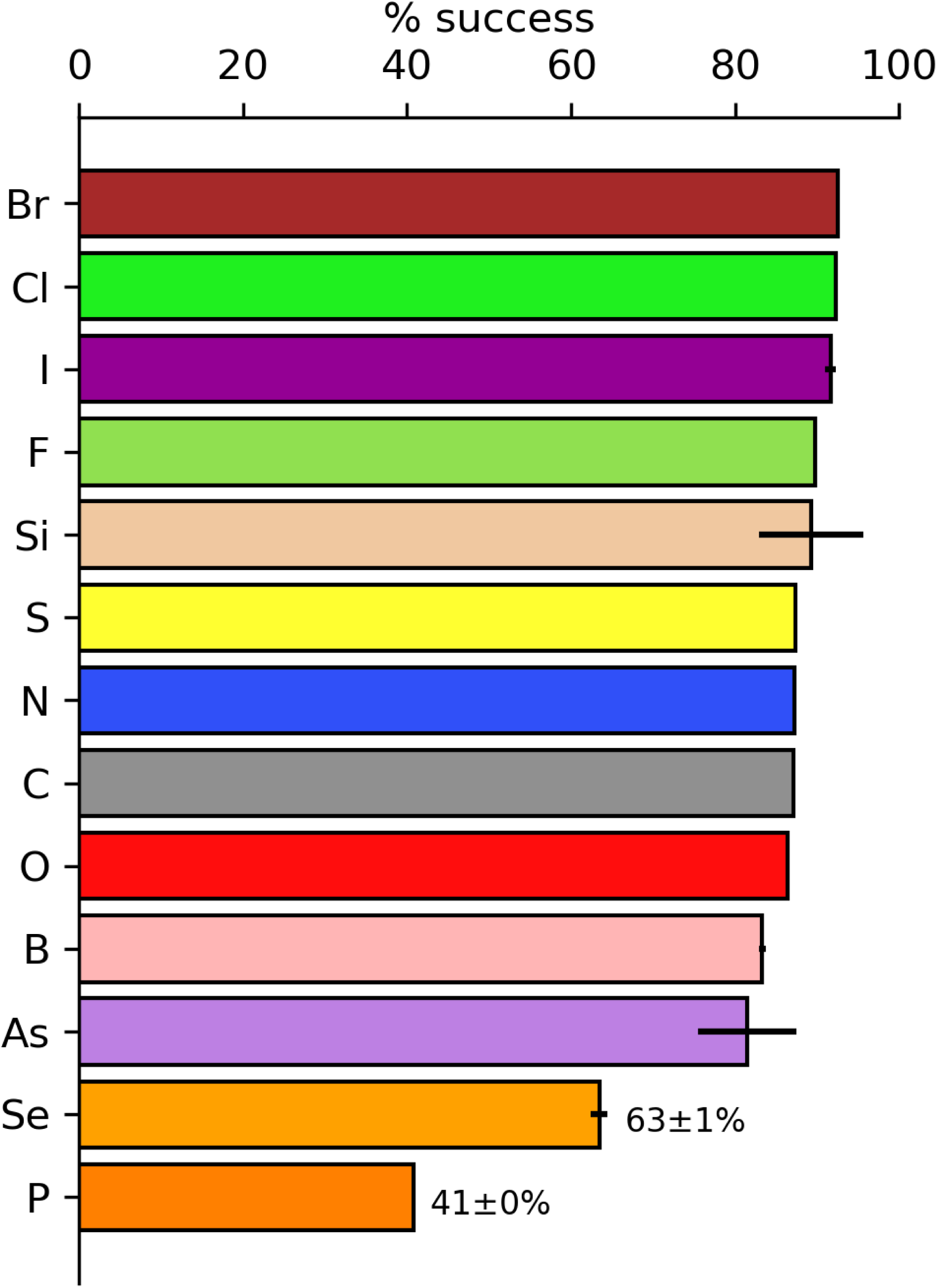
Success rate of molecules containing a given element using the CPK (CPK coloring, 2024) convention. Error bars are determined by the population of entities with the given element resulting in vanishingly small values. The values of success and error estimates are included for the lowest two elements.

The development of protonated specific restraints resulted in nearly 6k entries as shown in the last column of Table 1. Table 2 shows the number of validation classifications of the protonated and unprotonated states. The last column is the ligands that failed validation for the unprotonated ligand. When the corresponding protonated ligand was calculated, 149 were classified as “perfect”, 15 as “grand” and 405 as “ok”. The likely reason that most of the protonated ligands only passed as “ok” is because the unprotonated ligands were just outside the six Z-score cutoff. The largest number – 1835 – of ligands were classified as “grand” for the unprotonated entities and were upgraded for the protonated calculations to “perfect”. Clearly, the geometries are sensitive to the protonation state but there does not appear to be a clear trend. The number of entries that passed validation only for protonated geometries was 569 (last column sum).

**Table 2.** Number of entry classifications for the unprotonated geometries that validated into each classification for the protonated geometries.

|  | unprotonated |  |  |  |
| --- | --- | --- | --- | --- |
| protonated | perfect | grand | ok | fail |
| perfect | 313 | 1835 | 519 | 149 |
| grand | 753 | 445 | 55 | 15 |
| ok | 77 | 496 | 1017 | 405 |

As mentioned above, each of the components that were subject to force field generation were then fully minimized to the nearest local minimum in AmberTools. Figure 5 shows how much (as measured by r.m.s.d.) the MM-optimized structures deviated from the starting QM-optimized structures. Nearly 60% of the components changed less than 0.5Å upon MM minimization while 85% changed less than 1Å. Many of the molecules that changed more than this contain one or more rotatable bonds, with the MM optimum being in a different rotameric state. These cases do not lead us to believe the QM minimized geometry is necessarily inaccurate. The minimized structures and optimization details are also in the library, allowing users to assess the behavior of any particular component of interest.

**Figure 5.**
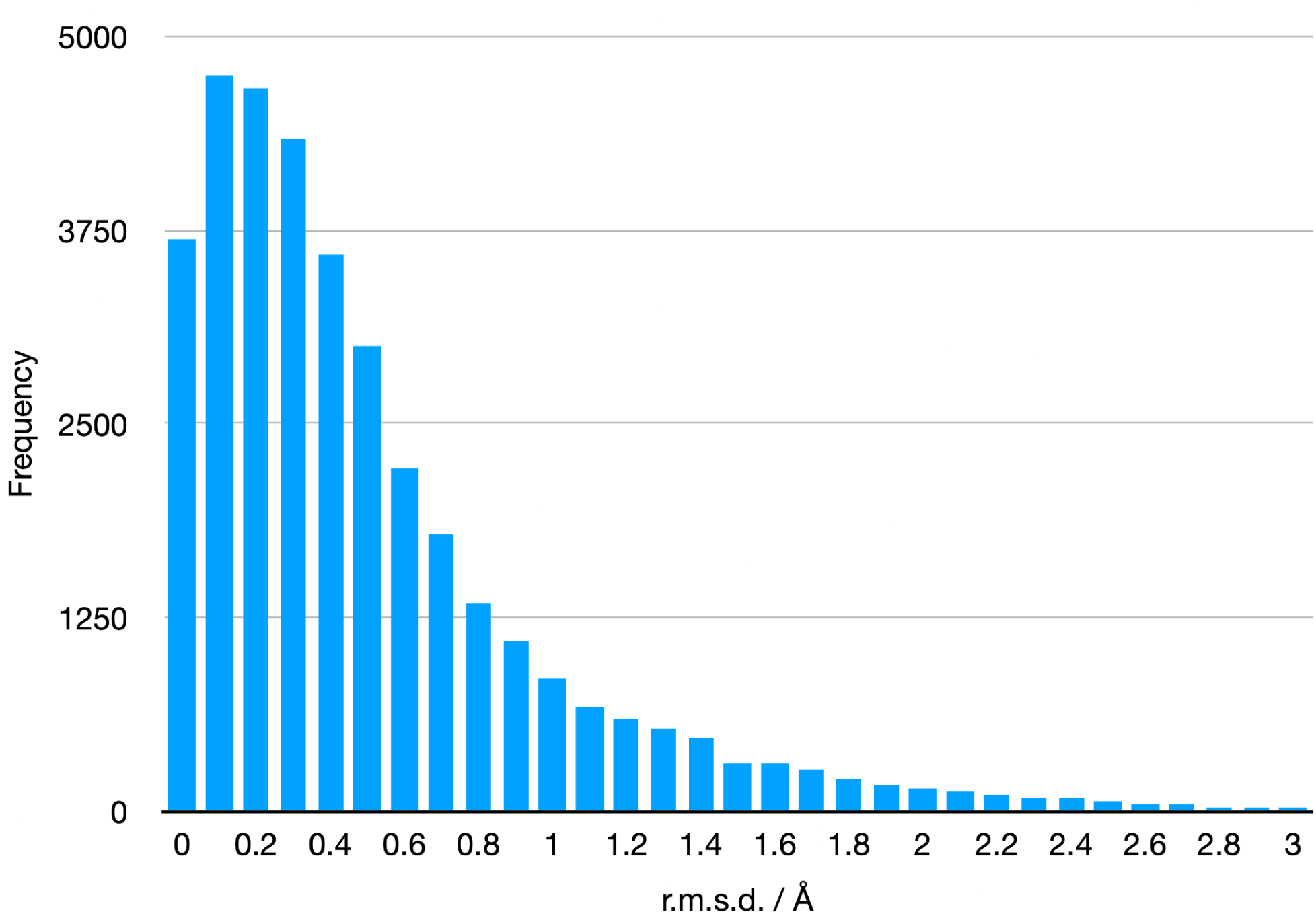
Differences between the starting and finishing geometry in r.m.s.d. of the Amber force field parameter files.

A similar validation was performed using the geometry minimisation program in Phenix (phenix.geometry_minimization). For each ligand, a geometry minimisation was performed starting with the ideal geometry from the restraints file. This procedure tests whether the restraints generated from the ideal geometry maintain that ideal geometry. The r.m.s.d. values between the starting and final geometry are shown in Figure 6. Because the geometry minimisation algorithm has a far simpler non-bonded interaction – only a repulsive potential – the r.m.s.d. distribution is skewed more towards zero than the MM force field minimisations. In fact, 87% are less than 0.5Å and 98.2% are less than 1.0Å. Rotatable bonds contributed to the larger r.m.s.d. values – for example, CCD ID: Fi3^2^ – but larger molecules tend to have larger r.m.s.d. values even when there are small deviations from the ideal restraints values that leverage the global measurement – for example, CCD ID: 3ZH.

**Figure 6.**
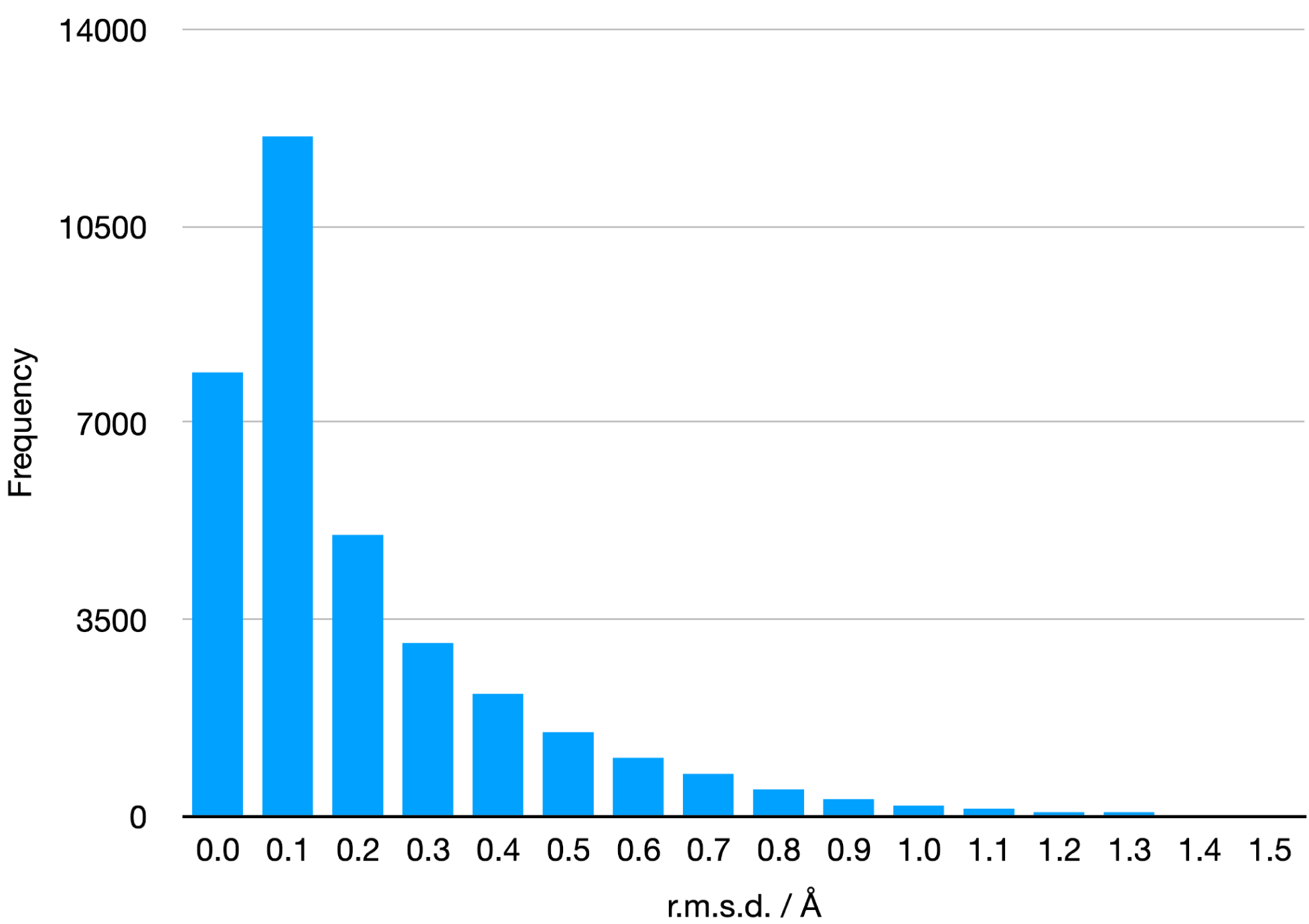
Differences between the starting and finishing geometry in r.m.s.d. of the restraints files.

The semiempirical QM method PM6-D3H4 was also assessed for suitability due to its ease of use (distributed with Phenix) and computer efficiency (Table S3). The general observation is that the PM6 geometries have just over 26k validated geometries compared to the PBEh-3c with nearly 33k. Furthermore, the majority of the successes are in the “ok” category compared to “perfect”. We conclude that while this level of QM has a lower success rate, it is highly computer efficient.

## Discussion

To improve on future iterations of this effort, we investigated entities not added to the GeoStd. First, the analysis showed that a significant number of models in the PDB have unknown entities designated with UNX (817) and UNL (550). Such molecules cannot be processed. Many of the next most populous entities are nucleotides including DPo – diphosphate – and metal clusters including CUA – dinuclear copper ion. Metal clusters can be determined via other methods but an investigation into phosphate moieties is warranted as one third of the entities that failed validation contained phosphorus. The development of an effective procedure for nucleic acids and nucleotides would greatly reduce the outstanding restraints as well as increase understanding of the chemistry of the phosphate moiety. The “missing” ligand restraints translates to less than 8% of PDB entries needing restraints beyond the GeoStd. This drops to 2.3% when ignoring RNA/DNA and metal entities – both of which are outside the scope of this work. It is likely that these are ligands of interest and, as such, deserve increased attention. It also highlights the need for deposition of restraints into the PDB for subsequent download.

We note that QM geometry minimisations work well for molecules with solely covalent bonds covering the majority of druglike candidates. Metals were excluded from this study because QM methods need additional attention including specification of electron configurations and higher levels of method and/or basis sets. Although this study was focused on covalent molecules, we did some tests with metals, which showed that even for metal containing ligands without electron configuration issues, Mg and Ca, the validation success was 38% and 20%, respectively.

The integration of Mopac (Moussa & Stewart, 2024) into the Phenix distribution has streamlined the use of semi-empirical QM for restraints generation. It is a complementary method to the use of Mogul for restraints generation. Furthermore, an interface with the Orca software package allows the use of the same methods verified in this work. Restraints for novel entities with druglike characteristics can be generated with confidence.

After generating restraints, it is important to validate the ligand geometry as the protein refinement progresses. We note that one can submit the model to the PDB for a validation report directly from the Phenix GUI. This is particularly important if the ligand is important to the results and reduces the possibility of surprises when the model and data are deposited.

## Conclusion

A procedure for generating and validating geometric restraints for ligand entities in the PDB Chemical Component Dictionary/Library (CCD) was developed and tested on over 42k entities. Each successfully validated set of restraints was added to the GeoStd – a freely available library of restraints in two formats (CIF) and Amber force-field files that can be downloaded from https://github.com/phenix-project/geostd. We note that this is also distributed with the Phenix software. Files relating to a particular CCD entry can be found at https://phenix-online.org/phenix_data/geostd/ by navigating to the appropriate directory.

The resulting restraint and molecular mechanics files can be used directly for crystallographic refinement or molecular dynamics simulations. The validation effort also illustrates the strengths and limitations of our current automated procedures for handling large numbers of chemical entities, such as small molecules and modified residues. Importantly, the restraints do not rely on the experimental geometry in the CSD directly. Using a validation method (in this case Mogul) to guide the refinement has challenges and can lead to a feedback loop that reinforces the possibly flawed metric. An independent method for generationing restraints is a positive step to independently verify the obtained geometries. By design the restraints generation and validation procedures discussed in this work can be repeated periodically as entries are added to the CCD. Furthermore, higher levels of QM can be investigated to address issues with difficult moieties.

## Availability

The Geometry Standard (GeoStd – pronounced Geo Standard) restraints library is available in the Phenix suite distribution (versions 1.21.2 and later) and at https://github.com/phenix-project/geostd. Individual entries may be accessed at https://phenix-online.org/phenix_data/geostd/.

## Acknowledgements

We thank the NIH (grant R24GM141254) and the Phenix Industrial Consortium for support of the Phenix project. This work was supported in part by the US Department of Energy under Contract No. DEAC02-05CH11231. The Molecular Sciences Software Institute is supported by NSF Grant No. CHE-2136142.

## Supplementary

**Table S1:**

| Size | PERFECT | PERFECT<br>(side<br>chain) | GRAND | GRAND<br>(side<br>chain) | OK | OK<br>(reasonabl<br>e std) | FAIL | Total<br>success | Total<br>Calculated |
| --- | --- | --- | --- | --- | --- | --- | --- | --- | --- |
| 10 | 699 | 12 | 738 | 67 | 136 | 115 | 630 | 1767 | 2397 |
| 20 | 3725 | 36 | 3387 | 478 | 1353 | 1113 | 3899 | 10092 | 13991 |
| 30 | 3797 | 8 | 3388 | 134 | 2422 | 2129 | 5626 | 11878 | 17504 |
| 40 | 2033 | 6 | 1887 | 57 | 2032 | 1808 | 3273 | 7823 | 11096 |
| 50 | 556 | 0 | 481 | 42 | 733 | 620 | 1190 | 2432 | 3622 |
| 60 | 158 | 2 | 120 | 13 | 202 | 201 | 532 | 696 | 1228 |
| 70 | 58 | 1 | 28 | 1 | 55 | 98 | 258 | 241 | 499 |
| 80 | 13 | 0 | 7 | 1 | 28 | 23 | 78 | 72 | 150 |
| 90 | 5 | 0 | 8 | 1 | 14 | 12 | 31 | 40 | 71 |
| 100 | 1 | 0 | 3 | 0 | 6 | 7 | 20 | 17 | 37 |
| 110 | 0 | 0 | 1 | 0 | 2 | 1 | 9 | 4 | 13 |
| 120 | 0 | 0 | 0 | 0 | 1 | 3 | 6 | 4 | 10 |

**Table S2:**
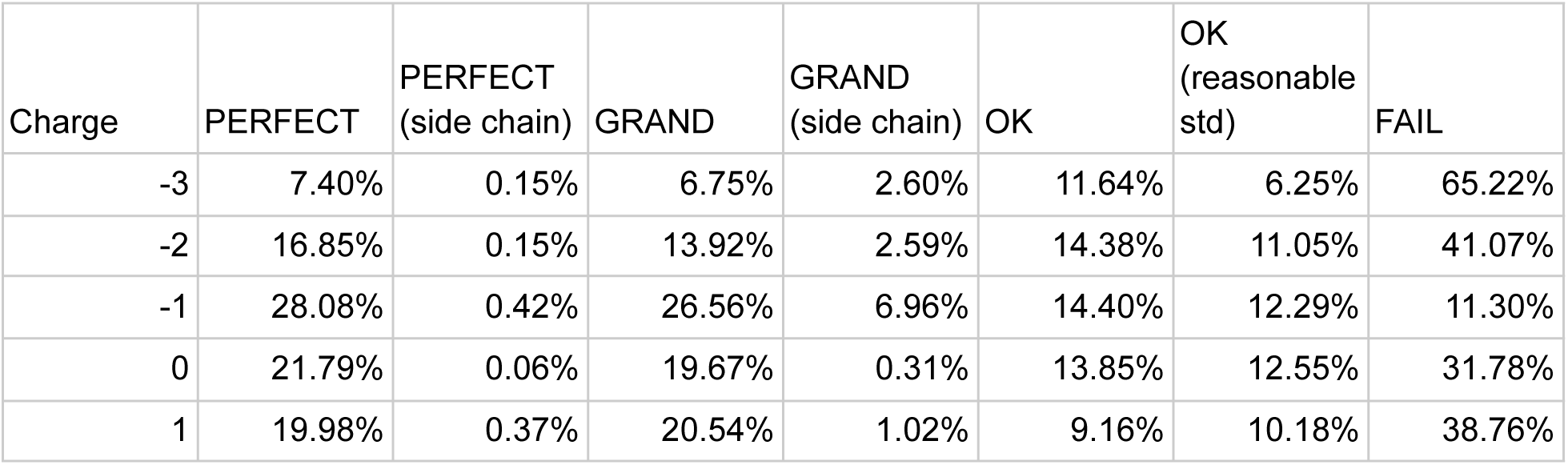

**Table S3:**

| Method | Mogul answer |  |  |
| --- | --- | --- | --- |
| PM6-D3H4 | GRAND |  | 1352 |
| PM6-D3H4 | GRAND | side chain | 166 |
| PM6-D3H4 | OK |  | 8501 |
| PM6-D3H4 | OK | reasonable std | 10347 |
| PM6-D3H4 | OK | side chain | 346 |
| PM6-D3H4 | PERFECT |  | 4346 |
| PM6-D3H4 | PERFECT | side chain | 28 |
|  |  | TOTAL | 25086 |

Original GeoStd Contents
ATOMN 5 [PSU]
ATOMP 456
ATOMS 9 [BMA, FUC, NGA, NDG, NAG, XLS, GAL, SIA, MAN]
HETAC 2 [NCO, IRI]
HETAI 4
HETAIN 55
HETAS 1
Total 532

## Footnotes

1 The CCD has dinuclear metal pairs as ions. Furthermore, Cl is a non-metal ion.

2 PDB and CCD ID codes are written using the “human readable” format as described in “Human Readable PDB Codes” — Moriarty, N.W. (2015). Comput. Crystallogr. Newsl. 6, 26.

